# DNA Release from Complex Plant Tissue using Focused Ultrasound Extraction (FUSE)

**DOI:** 10.1101/2022.06.23.497388

**Authors:** Alexia Stettinius, Hal Holmes, Qian Zhang, Isabelle Mehochko, Misa Winters, Ruby Hutchison, Adam Maxwell, Jason Holliday, Eli Vlaisavljevich

**Affiliations:** Department of Biomedical Engineering and Mechanics, Virginia Polytechnic Institute and State University, Blacksburg, VA, USA; Conservation X Labs, Seattle, WA, USA; Department of Forest Resources and Environmental Conservation, Virginia Polytechnic Institute and State University, Blacksburg, VA, USA; Department of Urology, University of Washington, Seattle, WA, USA

## Abstract

Sample preparation in genomics is a critical step that is often overlooked in molecular workflows and impacts the success of downstream genetic applications. This study explores the use of a recently developed focused ultrasound extraction (FUSE) technique to enable the rapid release of DNA from plant tissues for genetic analysis. FUSE generates a dense acoustic cavitation bubble cloud that pulverizes targeted tissue into acellular debris. This technique was applied to leaf samples of American chestnut (*Castanea dentata*), tulip poplar (*Liriodendron tulipifera*), red maple (*Acer rubrum*), and chestnut oak (*Quercus montana*). We observed that FUSE can extract high quantities of DNA in 9-15 minutes, compared to the 30 minutes required for conventional DNA extraction. FUSE extracted DNA quantities of 24.33 ± 6.51 ng/mg and 35.32 ± 9.21 ng/mg from American chestnut and red maple, respectively, while conventional methods yielded 6.22 ± 0.87 ng/mg and 11.51 ± 1.95 ng/mg, respectively. The quality of the DNA released by FUSE allowed for successful amplification and next-generation sequencing. These results indicate that FUSE can improve DNA extraction efficiency for leaf tissues. Continued development of this technology aims to adapt to field-deployable systems to increase the cataloging of genetic biodiversity, particularly in low-resource biodiversity hotspots.

## 1. Introduction

Over the past two decades, developments in genome sequencing technologies have enabled researchers to provide an unprecedented scope and depth of genetic information. Emerging technologies have equipped researchers with the tools to perform DNA and RNA sequencing in the field [1], which could allow for new genetics research to be carried out by non-scientists in a variety of settings that had not previously been feasible [2–4]. Despite technological advancements, novel sequencing platforms cannot be applied to all sample types, including many plant species, due to the poor representation of plant genomes in genetic databases. For example, of the nearly 400,000 unique plant species estimated to exist, only 600 have nearly complete genome coverage and assembly [5]. A recent review surveying commonly referenced databases found that only 17.7% of plants had broad genetic coverage, 80% of plant species had limited data availability, and some had no information other than their taxonomic names reported [6]. With such little coverage of plant taxa, it is likely that many opportunities for new uses of undiscovered traits unique to species have gone unnoticed, and with extinction rates rising, we may lose some of these opportunities forever. Sequencing plant genomes is also essential for utilizing genetic resources in breeding programs [7], conserving plant species [8, 9], understanding their role in ecosystem function [6, 10, 11], and phylogenetic studies [12]. Therefore, continued expansion of plant genetic databases is essential to spur discovery, drive innovation, and protect crucial resources. To accomplish this, tools for more efficient DNA extraction are necessary. Despite recent advancements in genome sequencing, DNA extraction technologies have lagged behind, particularly for complex plant tissues where the cell walls and membranes need to be broken down without significantly degrading the genetic material.

Therefore, sample preparation and DNA extraction remain a painful barrier that prevents rapid and inexpensive sequencing of plant genomes.

All genetic testing platforms require the input of purified genetic material. Consequently, a robust DNA extraction protocol that yields DNA of sufficient concentration and purity is essential for success in subsequent genotyping and sequencing applications. In plants, the release of viable DNA is hindered by tough tissue matrices that are resistant to mechanical breakdown, the presence of polysaccharide-rich cell walls, and many inhibitory compounds such as polyphenolic metabolites [4, 13, 14]. To combat these challenges, manual tissue pulverization with benchtop tools, such as a mixer mill, or a mortar and pestle under liquid nitrogen, is used in conjunction with plant cell lysis and purification protocols. Plant DNA extraction is often cumbersome, and despite specialized tissue breakdown strategies, releasing DNA suitable for genomic analysis is challenging for many sample types. Additionally, current DNA extraction techniques require an advanced laboratory [15, 16]. With the rise of point-of-contact genetic testing, the ability to translate DNA extraction protocols to the field is becoming increasingly important [3, 17]. For plants, the simplification of DNA extraction could be pivotal in conservation efforts where researchers must rapidly and inexpensively prepare samples from biodiversity hotspots, which are often remote and far removed from centralized laboratories [3, 18, 19].

Our group has recently developed a new technology capable of accelerating and simplifying the DNA extraction workflow, termed focused ultrasound extraction (FUSE), to address sample preparation and DNA extraction challenges. FUSE has previously demonstrated its capacity to rapidly release DNA from Atlantic salmon muscle tissue samples with intense cavitation clouds generated by focused ultrasonic transducers [20]. This technology employs dense acoustic cavitation bubble clouds similar to those used in histotripsy, a non-invasive focused ultrasound therapy currently being developed for medical applications [21, 22]. During FUSE, the rapid expansion and violent collapse of the cavitation microbubbles induce high stress on the target tissue, which causes mechanical disintegration and results in an acellular tissue lysate [23, 24]. The tissue lysate is then collected, and the released DNA is purified for downstream analyses. This process differs from conventional extraction methods that require samples to be pulverized by hand using a mortar and pestle or automated homogenizer under liquid nitrogen and require elongated incubation periods varying from 10 minutes to 1 hour, depending on the plant tissue, before DNA collection and purification (Figure 1).

**Figure 1:**
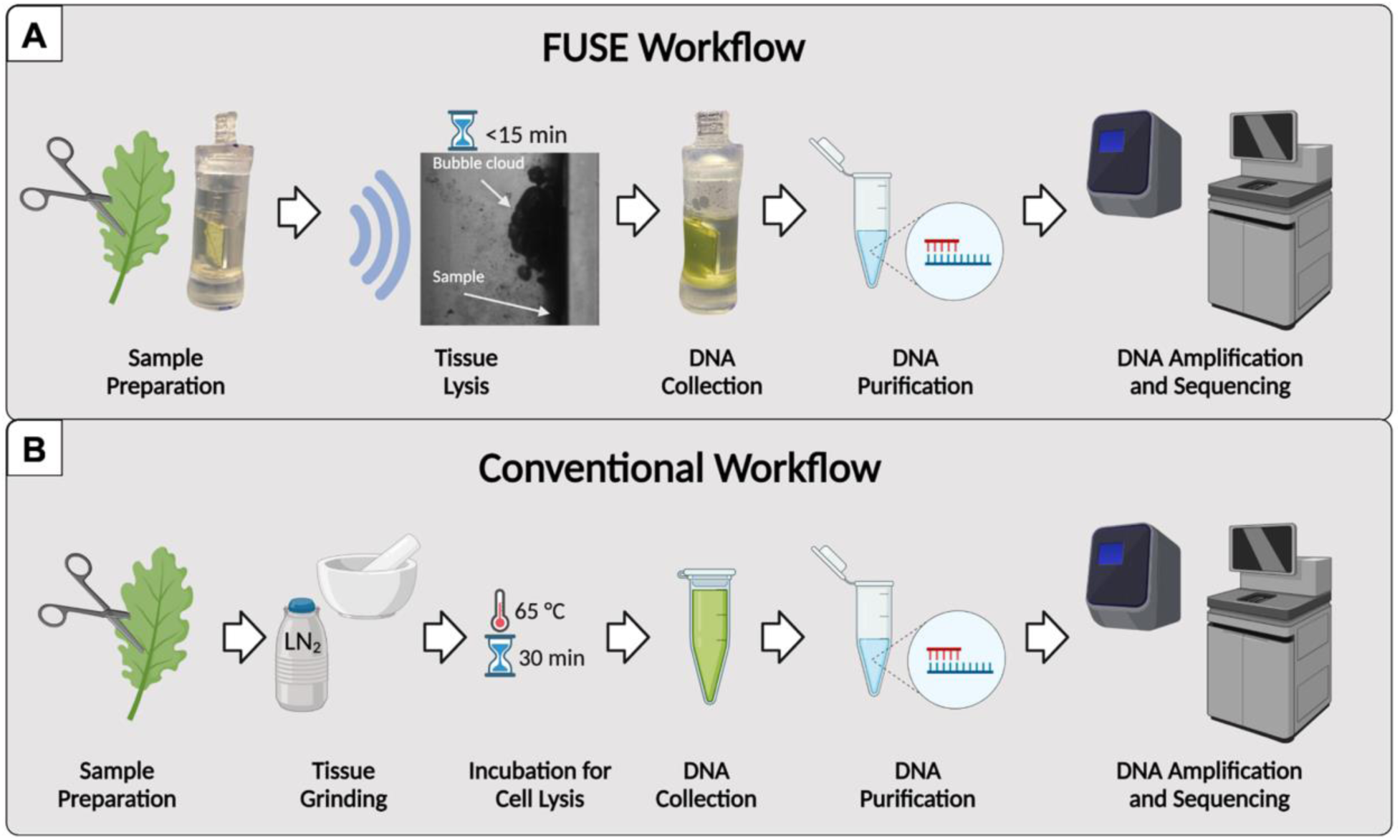
DNA extraction workflow. The process begins with sample preparation for FUSE and conventional DNA extraction protocols. In both cases, leaf samples are trimmed and massed before tissue processing. For FUSE, the prepared sample is aligned with the cavitation bubble cloud, and the tissue is sonicated in 15 minutes or less, eliminating the need for incubation. In both protocols, the DNA is then collected and purified. In the conventional extraction protocol, tissue processing involves manual grinding under liquid nitrogen and a 30-minute minimum incubation period. Following purification, the samples are prepared for amplification and sequencing.

Here, we test the efficacy of FUSE with plant tissue by 1) determining the feasibility of leaf tissue breakdown, 2) measuring the DNA yield, 3) amplifying the released DNA with PCR to verify DNA quality, and 4) sequencing the resultant DNA libraries to validate the utility of FUSE in whole-genome sequencing applications. While our ultimate goal is to adapt FUSE to low-cost and field-deployable systems to enable rapid sample processing and DNA extraction from various sample types, here we address the feasibility of FUSE for DNA release from leaf tissue in a laboratory setting using prototype transducers and custom acoustically transparent sample holders. We hypothesize that FUSE can pulverize leaf tissue and yield significant quantities of DNA with a quality suitable for PCR amplification and next-generation sequencing. If successful, FUSE could streamline plant DNA extraction workflows to improve standard laboratory practices. Further technology development could allow the miniaturization of the FUSE system to bring this technology to the field to expand the scope of opportunities.

## 2. Materials and Methods

To demonstrate the ability of FUSE to rapidly provide amplifiable DNA fragments for genetic analysis, we used frozen leaf samples collected from American chestnut (*C. dentata*), tulip poplar (*L. tulipifera*), red maple (*A. rubrum*), and chestnut oak (*Q. montana*) trees that were stored at -20 °C before use. These species were selected because they were locally available and represented a wide range of angiosperm taxonomic diversity that may correspond to variation in physical properties and secondary metabolite composition. The tissue samples were prepared and processed under the following experimental conditions:

### 2.1. FUSE Pulse Generation

A custom 32-element 500 kHz array transducer with a geometric focus of 75 mm, an aperture size of 150 mm, and an effective f-number of 0.58 was used for all experiments in this study [25]. A custom high-voltage pulser was used to drive the transducer and generate a short single cycle ultrasound pulse. The pulser was connected to a field-programmable gate array (FPGA) board (Altera DE0-Nano Terasic Technology, Dover, DE, USA), which was explicitly programmed for histotripsy therapy pulsing. A custom-built fiber-optic probe hydrophone (FOPH) [26] was used to measure the acoustic output pressure of the transducers. The FOPH was cross-calibrated at low-pressure values using a reference hydrophone (Onda HNR-0500) to ensure accurate pressures were measured from the FOPH. The lateral and axial full width half maximum (FWHM) dimensions at the geometric focus of the transducer were measured to be 2.3 mm and 7.1 mm, respectively. The acoustic pressures used for all experiments were measured in degassed water at the focal point of the transducer, which was identified using a 3D beam scan. The acoustic output could not be directly measured at higher pressure levels (p-> 16 MPa) due to cavitation at the fiber tip. These pressures were estimated by summating the output focal *p-* values from individual transducer elements. For all samples in this study, a pressure of ∼34 MPa was applied for FUSE processing.

### 2.2. Visualization of FUSE Tissue Disintegration

For all focused ultrasound experiments, high-speed optical imaging was done using a machine-vision camera (Blackfly S 3.2MP Mono USB3 Vision, FLIR Integrated Imaging Solutions, Richmond, BC, Canada) that was aligned with the focal zone of the transducer using a 100 mm F2.8 Macro lens (Tokina AT-X Pro, Kenko Tokina Co., LTD, Tokyo, Japan) and backlit by a custom-built pulsed LED strobe light capable of high-speed triggering with 1 µs exposures. As done in previous studies, the camera and the strobe light were triggered individually by the amplifier box, with the camera shutter opening at the time of pulse generation and the strobe acting as the shutter [27, 28]. The camera was triggered to capture one image every 50^th^ pulse. The exposures were centered at delay times of 6, 48.5, and 98.5 μs after the pulse arrived at the focus to allow visualization of bubble cloud formation, coalescence, and collapse.

### 2.3. Sample Preparation

Three leaves of American chestnut, tulip poplar, red maple, and chestnut oak were processed with FUSE and conventional extraction methods. Half of each leaf was used for FUSE, and the other half was used for conventional extraction (Figure S1). Three samples were acquired from each half. All samples processed with FUSE were prepared as 12 mm squares using a sterile scalpel blade. The mass of FUSE samples ranged from 10-30 mg, depending on the thickness of the sample.

#### 2.3.i FUSE Experimental Configuration

Leaf tissue samples were secured in a custom-designed sample holder positioned in the axial focus of the transducer, located between the camera and light source. The sample holder was designed to support a 12.5 mm x 12.5 mm x 1 mm sapphire glass window, the sample, and a polyethylene terephthalate glycol (PETG) square frame that secured the sample on the surface of the glass backing. The assembled sample holder was placed inside an optically transparent and acoustically permeable tube with an inner diameter of 9.525 mm and a wall thickness of 1.59 mm (Tygon PVC E-1000, McMaster-Carr, Douglasville, GA, USA). When the sample holder was placed in the tube, cylindrical appendages at the top and bottom of the sample holder created a controlled volume chamber for the DNA lysis buffer. The upper appendage featured a small circular opening for applying the DNA lysis buffer. A custom-built mount suspended the tube assembly in the water tank for tissue processing, and a stopper was designed to seal the other end of the tube. A robotic positioning system controlled by custom MATLAB scripts was used to align samples with the focus of the ultrasonic transducer (Figure 2).

**Figure 2:**
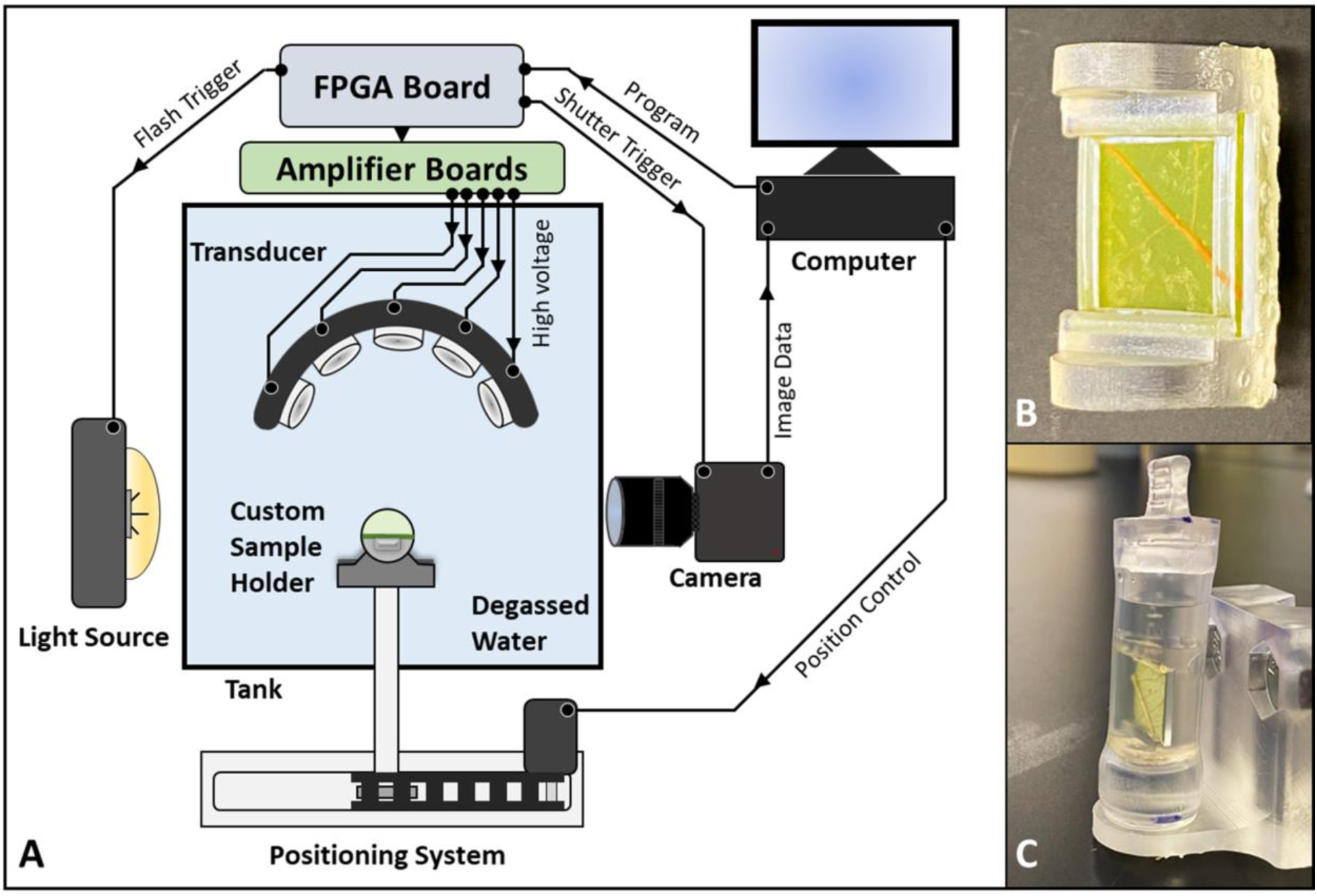
Experimental FUSE set-up. (A) Ultrasonic transducers are driven by an FPGA board and amplifier. High-speed imaging is performed using a strobe and camera controlled by signals from the FPGA board. Custom scripts are delivered to the FPGA board, and imaging data is recorded by a computer. A robotic positioning system, controlled by the computer using MATLAB, is used to align the sample in the focus of the transducer array. (B) A custom sample holder designed to support a sapphire glass backing, the leaf sample, and a PETG frame is used. (C) The sample holder assembly is housed in an acoustically permeable tube for DNA extraction experiments.

The configuration of the sample and sapphire glass backing in the focus of the transducer was chosen to maximize the efficiency of tissue sonication with FUSE. When ultrasonic pulses generate a cavitation bubble cloud near a rigid boundary, high-pressure collapse is expected to occur toward the surface of the boundary [29–31]. Figure 3 demonstrates this effect. As the time after pulse arrival increased, microbubble coalescence became more evident, and the concentration of bubbles near the sample surface increased. Sapphire glass was chosen because it is hydrodynamically strong and has a high acoustic impedance. The hydrodynamic strength of the sapphire glass provided an unyielding surface to support the sample when exposed to high-pressure fluid flow caused by cavitation. The high acoustic impedance increased the pressure near the boundary and induced the cavitation bubbles to grow larger and collapse more violently. Overall, this effect maximized the impact pressure felt by the sample. In preliminary experiments, FUSE was tested without including the sapphire glass backing and sample holder. With this configuration, the sample was free to move outside of the focal zone, which decreased the tissue disintegration efficiency of FUSE and caused inconsistencies in tissue breakdown success.

**Figure 3:**
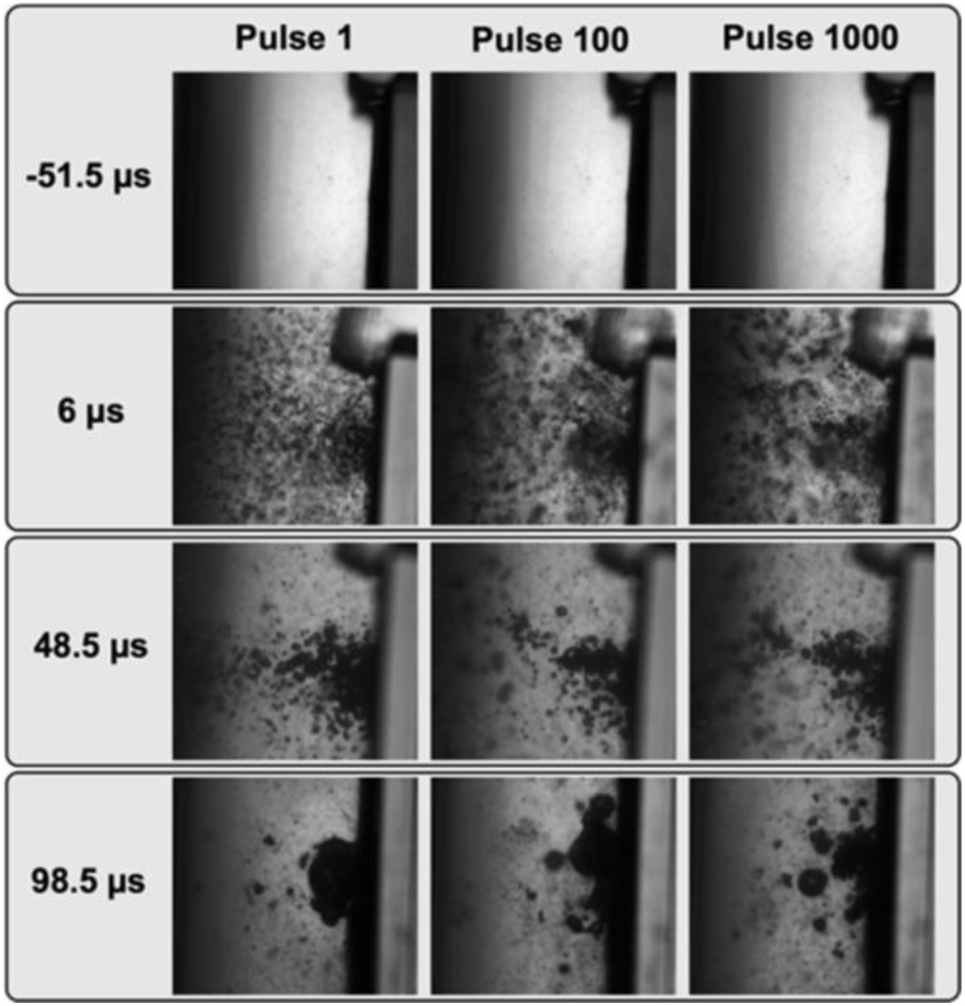
The cavitation bubble cloud collapses toward the surface of the leaf tissue. 51.5 μs before pulse arrival (row 1), the sample and sapphire glass backing are imaged. 6 μs after pulse arrival (row 2), the cavitation cloud is visible and contains many microbubbles that have not substantially expanded or coalesced. As time progresses (rows 3), the microbubbles begin to coalesce and are concentrated near the sample’s surface. 98.5 μs after pulse arrival (row 4), the microbubbles are near collapse and situated adjacent to the sample.

#### 2.3.ii. Control Sample Preparation and Tissue Lysis

Control samples were obtained by cutting the leaf tissue into 100 mg segments, and samples were disrupted by grinding with mortar and pestle under liquid nitrogen. 100 mg control samples were put in the mortar; liquid nitrogen was added to freeze the samples and cool the mortar, pestle, and spatula. To begin, grinding with the pestle was done slowly, and once the liquid nitrogen was mostly evaporated, more vigorous grinding was performed to reduce the tissue to a fine powder. The tissue powder was then transferred to a 1.5 mL centrifuge tube where 400 μL of 1% PVP-40 Buffer AP1 solution and 4 μL of RNase A were added (Qiagen DNeasy Plant Kit Qiagen Inc, Hilden, Germany). The tube was initially vortexed to homogenize the solution before incubation at 65 °C for 30 minutes with a short vortex every 5 minutes.

### 2.4 FUSE Tissue Disintegration

Leaf tissue samples were processed using single cycle ultrasound pulses delivered at a pressure of 34 MPa and a pulse repetition frequency (PRF) of 500 Hz. Measurements made with a fiber optic hydrophone determined that pressure loss was negligible (<1%) when pulses were delivered through the sample tube. Before tissue processing, the acoustic focus was directed at the center of the sample. To completely disintegrate the leaf tissue sample, MATLAB scripts controlling the positioning system were designed to move the sample in a spiral square pattern such that each point within the 100 mm^2^ disintegration zone was exposed to the focal bubble cloud for 0.5 s. Using this approach, a single scan of the applied pattern delivered 250 pulses per point, with multiple scans used for each sample to achieve sufficient tissue breakdown. To account for potential differences in the physical properties of the selected leaf species, the number of scans required for complete tissue disintegration was initially characterized for American chestnut, tulip poplar, red maple, and chestnut oak samples (n=3) backed with sapphire glass suspended in an open water bath. Images of the sample were taken after each scan. Each image was converted to grayscale, then to binary using the Otsu method [32]. The targeted tissue area was mapped as an ROI with dimensions of 10 x 10 mm to represent the area exposed to ultrasonic pulses. The disrupted tissue area inside and outside the ROI was quantified by counting the number of pixels using custom MATLAB scripts. Pixel counts were converted to tissue disintegration area. The significance of the area measurements was determined using an unpaired student’s t-test with unequal variance. Values less than 0.05 (p<0.05) were considered significant.

### 2.5 Purification Conditions

The robustness of the DNA extraction process was investigated through purification of the disintegrated tissue and quantification of the released DNA. DNA was extracted from samples of American chestnut, tulip poplar, red maple, and chestnut oak with FUSE (n = 9) and conventional extraction methods (n = 9) using a lysis buffer and purified using silica columns (Qiagen DNeasy Plant Kit Qiagen Inc, Hilden, Germany). All lysates were analyzed with a Qubit^TM^ 4 Fluorometer (ThermoFisher, Waltham, MA, USA) and a Nanodrop^TM^ One (ThermoFisher, Waltham, MA, USA) to determine the quantity and quality of DNA released with FUSE and control samples. DNA yield was reported as the quantity of DNA released per milligram of tissue to normalize input sample mass. For data acquired from Nanodrop^TM^ and Qubit^TM^ measurements, an unpaired student’s t-test with unequal variance was used, with values less than 0.05 (p<0.05) considered significant.

FUSE and control samples were purified with a lysis buffer containing 1% PVP-40 Buffer AP1 solution and RNase A. The lysis buffer used on the FUSE samples consisted of 1 mL of 1% PVP-40 Buffer AP1 solution and 8 μL of RNase A (0.8 mg), and samples were soaked for the duration of FUSE tissue sonication. The lysis buffer volume used for the controls varied depending on the quality of the leaf tissue sample, which was determined based on the color and age assessment. For older samples with a dark green or brown-green color, 1 mL of 1% PVP-40 Buffer AP1 solution and 8 μL of RNase A (0.8 mg) were used. For younger samples with a yellow-green color, 500 μL of 1% PVP-40 Buffer AP1 solution and 4 μL of RNase A (0.4 mg) were used. This was done because the leaves with a lower water content yielded a larger sample volume after grinding with mortar and pestle under liquid nitrogen and therefore required a greater buffer volume for proper cell lysis. Subsequent purification of FUSE and control samples was performed in silica columns using the standard protocol as recommended by the manufacturer (Qiagen DNeasy Plant Kit Qiagen Inc, Hilden, Germany).

### 2.6 PCR Amplification

To compare the two methods for downstream genotyping, American chestnut genomic DNA extracted by FUSE and conventional methods was subjected to a genotyping by sequencing (GBS) workflow that involved restriction digestion followed by ligation of sequencing adapters and PCR amplification [33]. American chestnut samples processed with FUSE and conventional methods were normalized to 55 ng, then digested with 1 μL of ApeKI (New England BioLabs, Ipswitch, MA, USA). This restriction enzyme recognizes a 5 bp degenerate sequence GCWGC, where W is an A or T [34]. For one of the American chestnut samples processed with FUSE, the quantity of eluted DNA did not reach 55 ng, so 36.2 ng of DNA was used in the digestion reaction. The resulting DNA fragments were ligated to Illumina-compatible adapters with 1.6 μL of T4 DNA ligase. P1 adapters contained a unique barcode region for each adapter immediately upstream of the ligated DNA fragment, and the P2 adapter was consistent for all samples. PCR was performed under the following conditions: 95 °C for 1 minute, followed by 18 cycles of 95 °C for 30 seconds, 63 °C for 20 seconds, and 68 °C for 30 seconds. Lastly, samples were brought to 68 °C for 5 minutes and kept at 4 °C. PCR primers contained complementary sequences for amplifying restriction fragments with ligated adapters [34]. Ligation and amplification were assessed by gel electrophoresis for all samples. Six samples, three processed with FUSE and three processed with conventional methods, were viewed using a 2100 Bioanalyzer instrument (2100 Bioanalyzer, Agilent, Santa Clara, CA, USA) to compare the fragment size distribution for individual PCR products yielded by FUSE and conventional methods. DNA samples were purified before and after PCR with the Monarch® PCR and DNA Clean-Up Kit (New England Biolabs, Ipswitch, MA, USA). Individual sample libraries were pooled, and fragments ranging from 250-550 bp were selected using BluePippin™ (Sage Science, Beverly, MA, USA). The resulting library was visualized using a 2100 Bioanalyzer instrument.

### 2.7 Sequencing Analysis

The American chestnut GBS libraries were sequenced with the NovaSeq 6000 instrument (Illumina, San Diego, CA, USA) in 2x150 bp paired-end mode at Duke University Center for Genomic and Computational Biology. Raw reads were filtered for quality and adapter contamination, then demultiplexed using STACKS software [35]. Filtered reads were aligned to v.1.1 of the *C. dentata* reference genome [36] using the Burrows-Wheeler Aligner (BWA) mem algorithm and subsequently converted to BAM format with SAMtools [33]. Heterozygous sites were called with the Genome Analysis Toolkit (GATK) HaplotypeCaller algorithm [37, 38], and these GVCFs were then merged using the GenotypeGVCFs function. Variants were flagged and removed as low quality if they had the following characteristics: low map quality (MQ < 40); high strand bias (FS > 40); differential map quality between reads supporting the reference and alternative alleles (MQRankSum < -12.5); bias between the reference and alternate alleles in the position of alleles within the reads (ReadPosRankSum < -8.0); and low depth of coverage (DP < 5). Coverage depth per sample was calculated using the SAMtools depth function. Statistical analysis of coverage depth was performed using a Wilcoxon rank-sum test with values less than 0.05 (p<0.05) considered significant.

## 3. Results & Discussion

### 3.1 FUSE Tissue Disintegration

The feasibility of FUSE for leaf tissue disintegration was examined with American chestnut, tulip poplar, red maple, and chestnut oak leaves by characterizing tissue breakdown after each FUSE scan. Images were captured after each scan to demonstrate the progression of tissue disintegration for each species (Figure 4). In all cases, the damaged tissue area increased with increasing scan number, but the number of scans required to achieve significant tissue breakdown differed among species. Tulip poplar leaves were the most vulnerable to breakdown, as they were the only species with notable tissue disintegration after one scan. The extent of tulip poplar tissue disruption increased with each scan. For the American chestnut and red maple samples, scans one and two generated minimal tissue breakdown, but after scan three, the area of tissue breakdown was more observable. For American chestnut, the tissue in the targeted region was completely disrupted after four scans. For red maple, the tissue breakdown continued to increase after scans four, five, and six. For American chestnut, tulip poplar, and red maple samples, tissue breakdown beyond the bounds of the targeted disintegration zone was observed. This effect was likely due to dispersed cavitation occurring outside the focal point of the converging pressure fronts. Surface inhomogeneities at solid-liquid interfaces result in the growth of cavitation nuclei that can induce cavitation at thresholds below the intrinsic threshold [39, 40]. Previous work has also shown that leaves are more susceptible to cavitation-induced tissue disruption when gas channels are present in the tissue [41, 42]. Therefore, it is possible that the surface architecture and distribution of gas channels within the tissue matrices created cavitation nucleation sites outside the targeted area. It is also possible that residual gas bubbles from preceding pulses diffused outside the focus and served as cavitation nuclei. This would induce cavitation below the intrinsic pressure threshold and expose a larger area of the leaves to cavitation [43]. Lastly, off-target leaf tissue disintegration could result from acoustic shielding, such that the residual bubbles in the acoustic focus increased the likelihood of acoustic scattering [44, 45]. The trends in tissue breakdown for chestnut oak differed from the other three species. Visible tissue breakdown was not observed until after the third scan, and tissue disintegration in the following scans did not progress as promptly as it did for the American chestnut, tulip poplar, and red maple samples. It is expected that variation in tissue breakdown across species was due to differences in physical properties, such as the water content and tissue strength.

**Figure 4:**
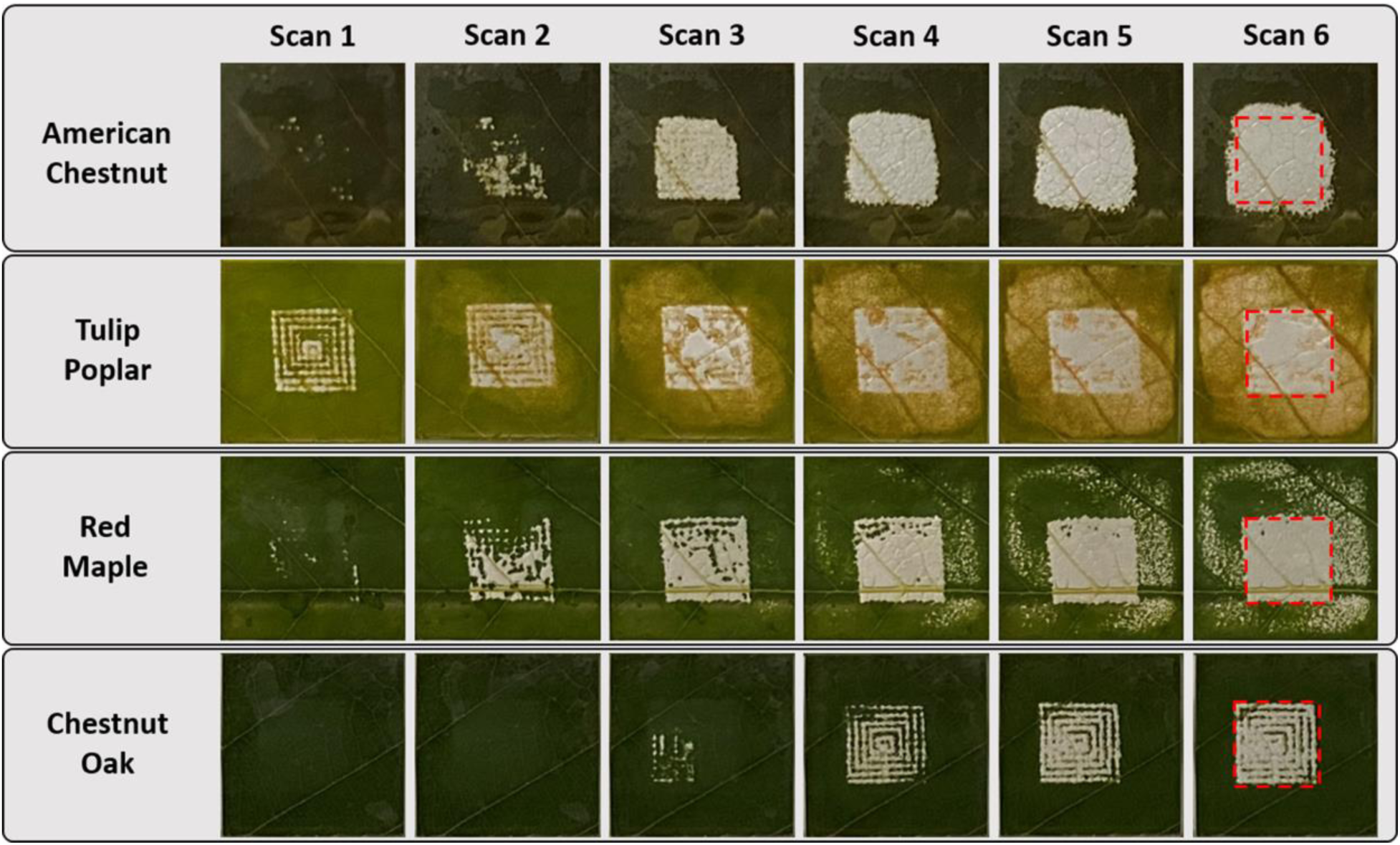
Leaf tissue disintegration increases after each FUSE scan. The red square in the top right identifies the targeted tissue region. Tissue breakdown beyond the target area is the result of peripheral cavitation damage. Image data suggests that the leaf species affect FUSE tissue disintegration efficiency.

The observed efficiency of FUSE tissue disintegration was assessed quantitatively by plotting the disintegration area inside the targeted region and the total disintegration area as a function of scan number for American chestnut, tulip poplar, red maple, and chestnut oak leaves (Figure 5). The initial breakdown occurred the most rapidly in tulip poplar leaves, as after scan one, 38.4 ± 11.4% of the targeted tissue region was disintegrated. In comparison, <10% of target tissue was disintegrated after scan one for American chestnut, red maple, and chestnut oak. The targeted tulip poplar tissue region was significantly processed after two scans (p<0.05 compared to zero scans), and >90% of the targeted area was processed after four scans. After six scans, a final disintegration area of 97.4 ± 0.34% was observed within the targeted region. The initial breakdown of American chestnut leaves did not occur as rapidly as tulip poplar, but American chestnut quickly approached complete tissue breakdown, reaching >90% of tissue breakdown inside the targeted area after three scans. American chestnut leaves were significantly processed after three scans (p<0.05 compared to zero scans), and six scans resulted in a disintegration area of 99.3 ± 0.75% inside the targeted region. Red maple leaves were more resistant to breakdown than tulip poplar and American chestnut, as four scans were required to achieve significant breakdown (p<0.05 compared to zero scans). Interestingly, the degree of breakdown for red maple in the targeted region never surpassed 90%. After six scans, the final disintegration area within the targeted region for red maple was 89.2 6.0%. The observed reduction in tissue breakdown efficiency could be due to the presence of the midrib, the central vein of the leaf. Previous work investigating the effects of ultrasonic cavitation on *Elodea* leaves found that when ultrasound was targeted at the midrib, there was a lack of cell disruption [42]. Although tulip poplar, American chestnut, and red maple samples achieved significant breakdown in less than six scans, increasing the scan number decreased the margin of error in the disintegration area. Therefore, six scans were used to process American chestnut, tulip poplar, and red maple samples to allow more consistent comparisons. Six scans resulted in a 9-minute tissue processing time and a total of 1,500 pulses per point. Chestnut oak was the most resistant to breakdown. After scan six, only 56.0 ± 22.9% of the targeted area was processed, so up to ten scans were applied to chestnut oak leaves. Chestnut oak samples required eight scans to achieve a significant breakdown of 76.6 ± 13.8% (p<0.05 compared to zero scans). Since increasing the scan number increased the area of disintegration and reduced the margin of error in the disintegration area, ten scans were used for chestnut oak tissue processing. Ten scans resulted in a 15-minute tissue processing time and 2,500 pulses per point. The final disintegration area within the targeted region for chestnut oak was 83.7 ± 10.6%.

**Figure 5:**
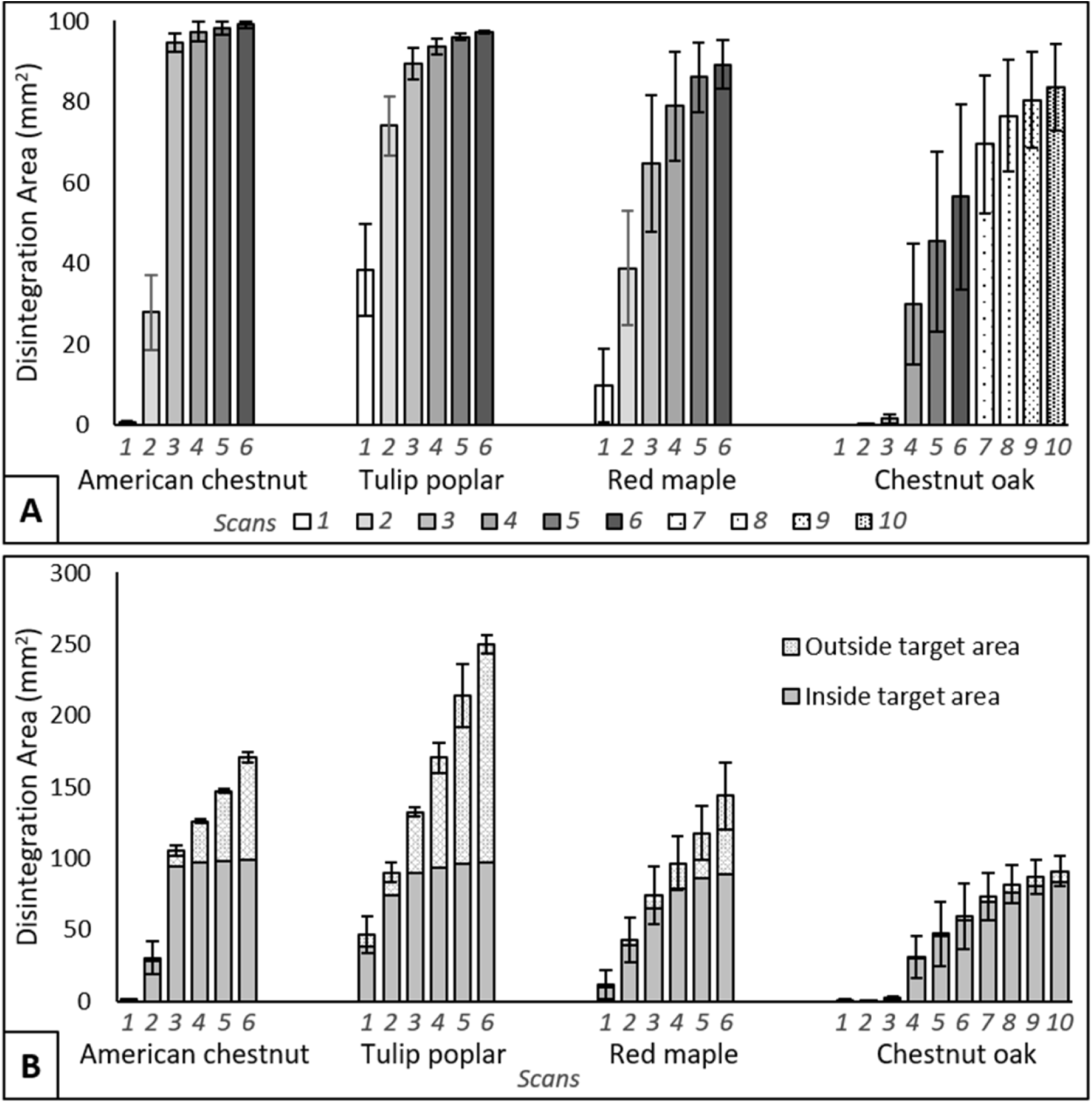
The leaf tissue disintegration area increases inside and outside the target area with the number of FUSE scans. (A) The disintegration area within the target area increases after each scan for all species. (B) The total disintegration area shows that tissue outside of the target area is also disintegrated by FUSE. Six scans are used for processing American chestnut, tulip poplar, and red maple samples. Chestnut oak samples required ten scans for processing due to a reduction in disintegration efficiency.

Leaf tissue breakdown outside of the targeted area was also quantified to examine the effects of dispersed cavitation. Trends in tissue breakdown outside of the targeted area were comparable to those observed within the targeted area, such that the tissue disintegration area increased with increasing scan number. Additionally, the extent of American chestnut, tulip poplar, and red maple tissue disintegration outside the targeted region was greater than chestnut oak. These results suggest that differences in the physical properties among leaf species also affect the extent of collateral tissue breakdown. However, collateral tissue breakdown is not of central importance for this study because the samples are restricted to the size of the targeted region in DNA extraction experiments.

### 3.2 DNA Extraction Feasibility

The determined number of FUSE scans required to disintegrate each leaf species was applied to the DNA extraction workflow to characterize the quantity of DNA released by FUSE compared to conventional methods, the control protocol in this study (Figure 6). Overall, FUSE was able to release greater quantities of DNA than conventional extraction methods in a fraction of the processing time. Notably, FUSE increased the DNA yield with less than half of the input sample mass required by the conventional protocol. The quantity of DNA released with FUSE and controls varied with species. The DNA yield provided by six FUSE scans was significantly greater than controls for American chestnut and red maple samples. Six FUSE scans released 24.3 ± 6.5 ng/mg from American chestnut and 35.3 ± 9.3 ng/mg from red maple samples, while controls yielded 6.2 ± 0.87 ng/mg and 11.5 ± 1.9 ng/mg, respectively. No significant differences were observed in the quantity of DNA released from tulip poplar samples between six FUSE scans, 32.6 ± 7.8 ng/mg, and controls, 28.4 ± 1.8 ng/mg. For chestnut oak leaves, 37.9 ± 5.9 ng/mg of DNA was provided by ten FUSE scans, 10.7 ± 1.7 ng/mg from six FUSE scans, and 17.2 ± 2.0 ng/mg from controls. The DNA yield provided by ten FUSE scans was significantly greater than six FUSE scans and controls for chestnut oak samples, showing that the capacity of FUSE to release DNA from tough tissues improves with an increased number of processing scans.

**Figure 6:**
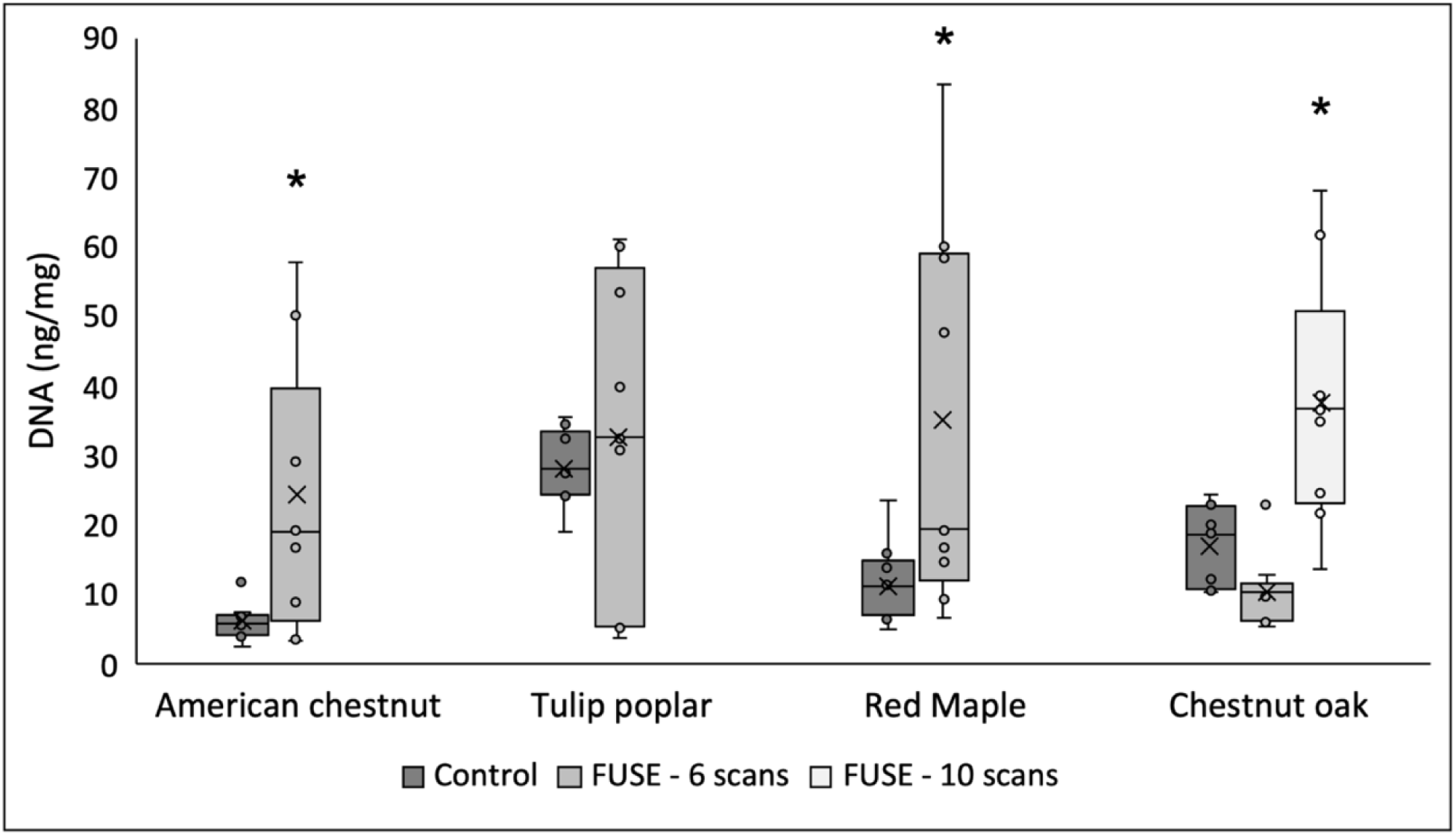
DNA extraction results show that FUSE releases DNA from leaf tissue. A significant increase in DNA release from American chestnut and red maple samples is observed when processed with 6 FUSE scans compared to controls. DNA release from chestnut oak samples is significantly higher when 10 FUSE scans are used for processing than controls. After 6 FUSE scans, DNA release from tulip poplar samples is comparable to controls. *Indicate significant (p<0.05) differences between FUSE and control samples.

**Table 1:**
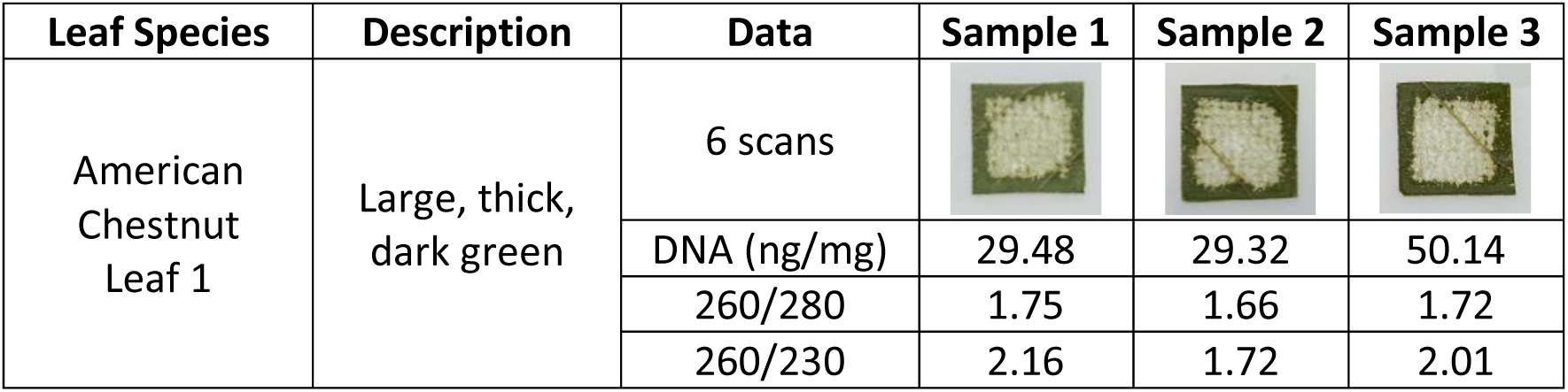
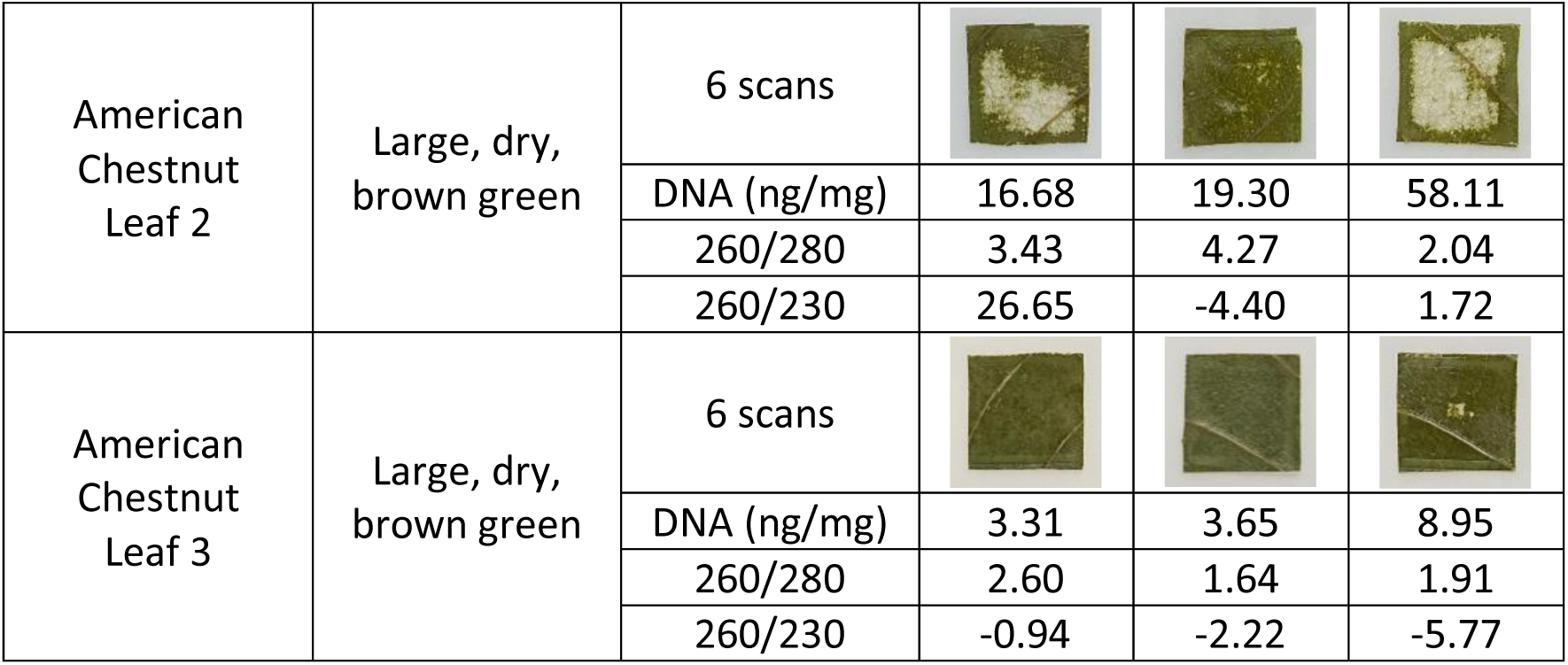
American chestnut tissue breakdown after FUSE processing demonstrates that greater tissue disintegration increases the concentration of DNA release. DNA quantification measurements are reported from Qubit^TM^ Fluorometer measurements, and 260/280 and 260/230 ratios are reported from Nanodrop^TM^ measurements.

| Leaf Species | Description | Data | Sample 1 | Sample 2 | Sample 3 |
| --- | --- | --- | --- | --- | --- |
| American Chestnut Leaf 1 | Large, thick, dark green | 6 scans |  |  |  |
|  |  | DNA (ng/mg) | 29.48 | 29.32 | 50.14 |
|  |  | 260/280 | 1.75 | 1.66 | 1.72 |
|  |  | 260/230 | 2.16 | 1.72 | 2.01 |
| American Chestnut Leaf 2 | Large, dry, brown green | 6 scans |  |  |  |
|  |  | DNA (ng/mg) | 16.68 | 19.30 | 58.11 |
|  |  | 260/280 | 3.43 | 4.27 | 2.04 |
|  |  | 260/230 | 26.65 | -4.40 | 1.72 |
| American Chestnut Leaf 3 | Large, dry, brown green | 6 scans |  |  |  |
|  |  | DNA (ng/mg) | 3.31 | 3.65 | 8.95 |
|  |  | 260/280 | 2.60 | 1.64 | 1.91 |
|  |  | 260/230 | -0.94 | -2.22 | -5.77 |

**Table 2:**
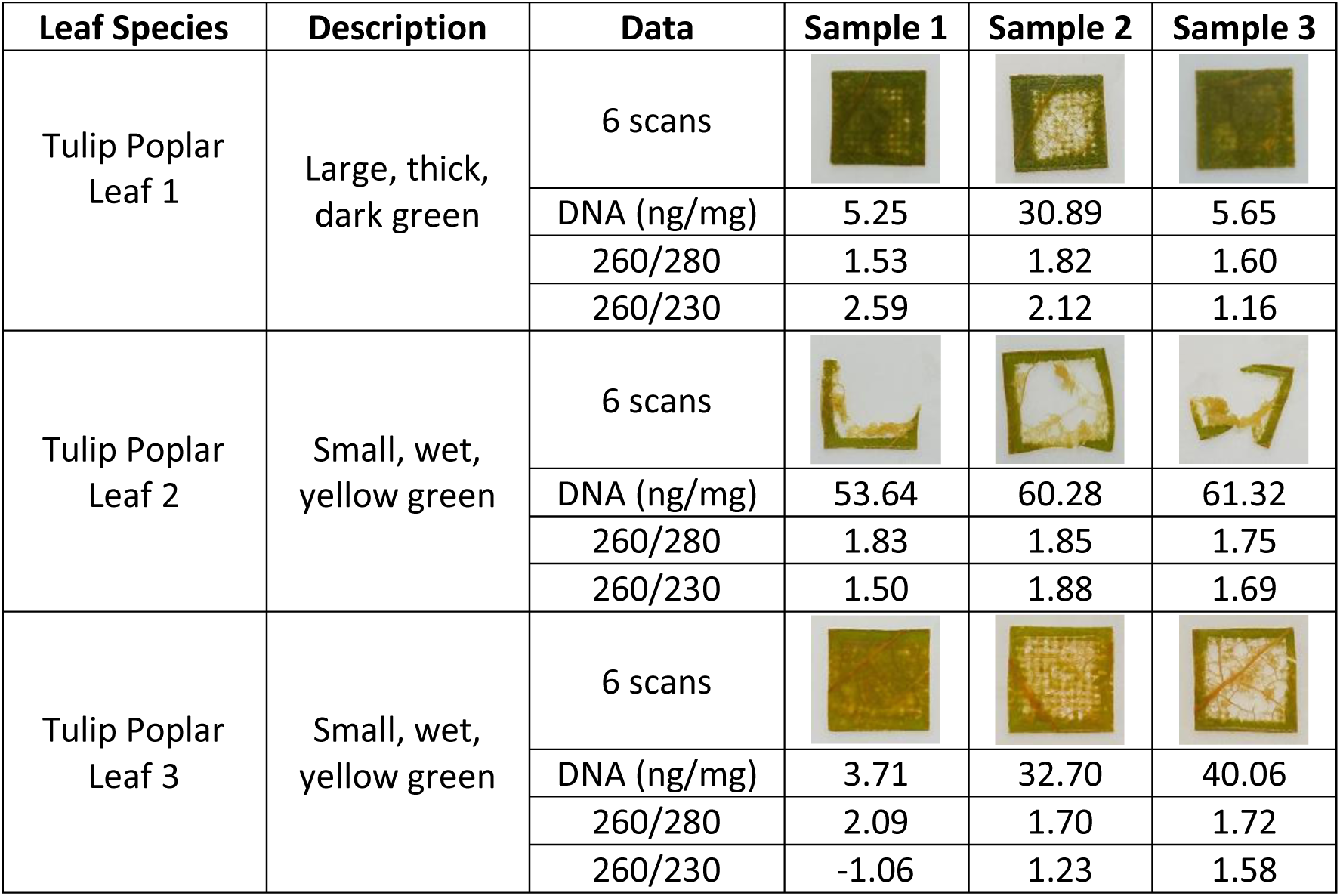
Tulip poplar tissue breakdown after FUSE processing demonstrates that greater tissue disintegration increases the concentration of DNA release. DNA quantification measurements are reported from Qubit^TM^ Fluorometer measurements, and 260/280 and 260/230 ratios are reported from Nanodrop^TM^ measurements.

| Leaf Species | Description | Data | Sample 1 | Sample 2 | Sample 3 |
| --- | --- | --- | --- | --- | --- |
| Tulip Poplar Leaf 1 | Large, thick, dark green | 6 scans |  |  |  |
|  |  | DNA (ng/mg) | 5.25 | 30.89 | 5.65 |
|  |  | 260/280 | 1.53 | 1.82 | 1.60 |
|  |  | 260/230 | 2.59 | 2.12 | 1.16 |
| Tulip Poplar Leaf 2 | Small, wet, yellow green | 6 scans |  |  |  |
|  |  | DNA (ng/mg) | 53.64 | 60.28 | 61.32 |
|  |  | 260/280 | 1.83 | 1.85 | 1.75 |
|  |  | 260/230 | 1.50 | 1.88 | 1.69 |
| Tulip Poplar Leaf 3 | Small, wet, yellow green | 6 scans |  |  |  |
|  |  | DNA (ng/mg) | 3.71 | 32.70 | 40.06 |
|  |  | 260/280 | 2.09 | 1.70 | 1.72 |
|  |  | 260/230 | -1.06 | 1.23 | 1.58 |

**Table 3:**
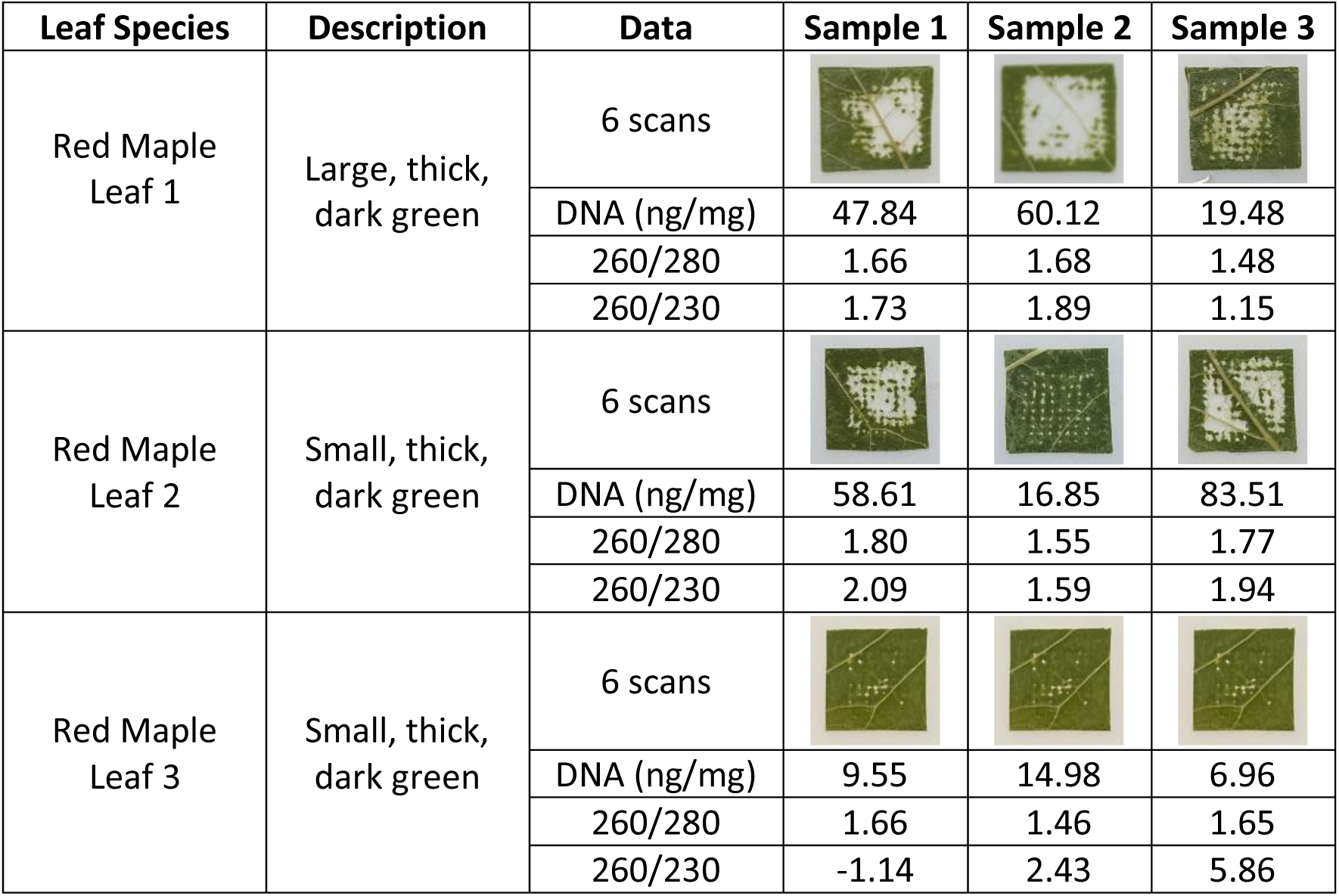
Red maple tissue breakdown after FUSE processing demonstrates that greater tissue disintegration increases the concentration of DNA release. DNA quantification measurements are reported from Qubit^TM^ Fluorometer measurements, and 260/280 and 260/230 ratios are reported from Nanodrop^TM^ measurements.

| Leaf Species | Description | Data | Sample 1 | Sample 2 | Sample 3 |
| --- | --- | --- | --- | --- | --- |
| Red Maple Leaf 1 | Large, thick, dark green | 6 scans |  |  |  |
|  |  | DNA (ng/mg) | 47.84 | 60.12 | 19.48 |
|  |  | 260/280 | 1.66 | 1.68 | 1.48 |
|  |  | 260/230 | 1.73 | 1.89 | 1.15 |
| Red Maple Leaf 2 | Small, thick, dark green | 6 scans |  |  |  |
|  |  | DNA (ng/mg) | 58.61 | 16.85 | 83.51 |
|  |  | 260/280 | 1.80 | 1.55 | 1.77 |
|  |  | 260/230 | 2.09 | 1.59 | 1.94 |
| Red Maple Leaf 3 | Small, thick, dark green | 6 scans |  |  |  |
|  |  | DNA (ng/mg) | 9.55 | 14.98 | 6.96 |
|  |  | 260/280 | 1.66 | 1.46 | 1.65 |
|  |  | 260/230 | -1.14 | 2.43 | 5.86 |

**Table 4:**
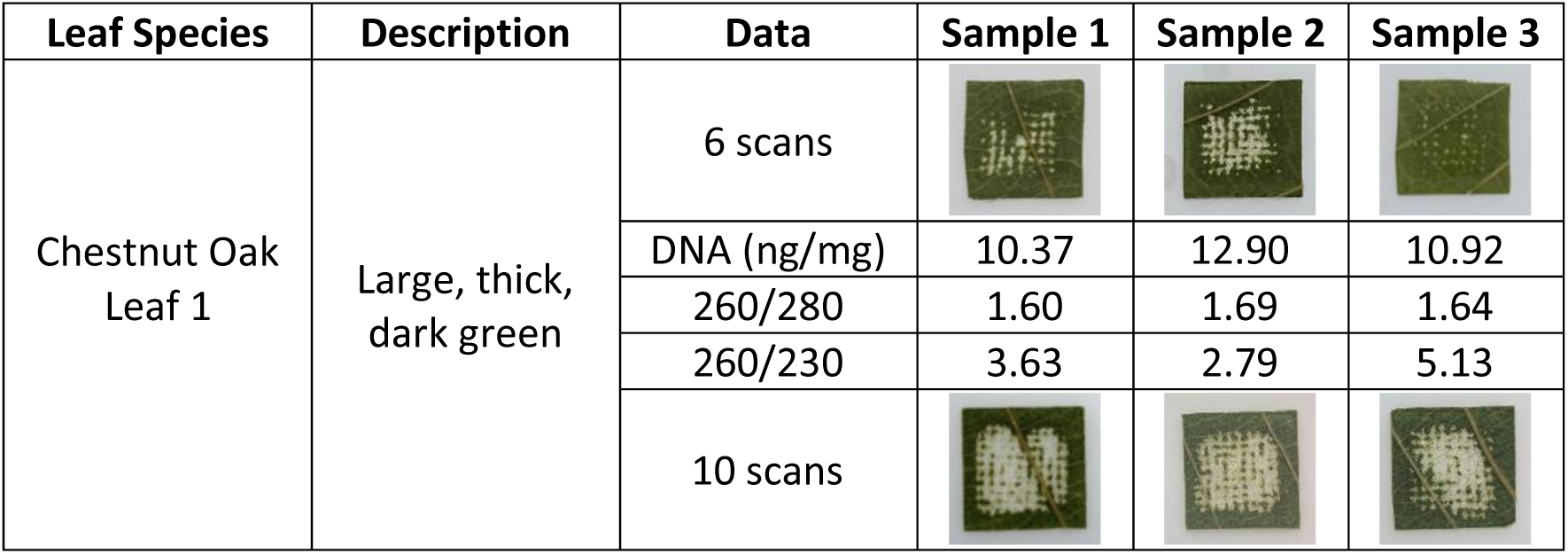
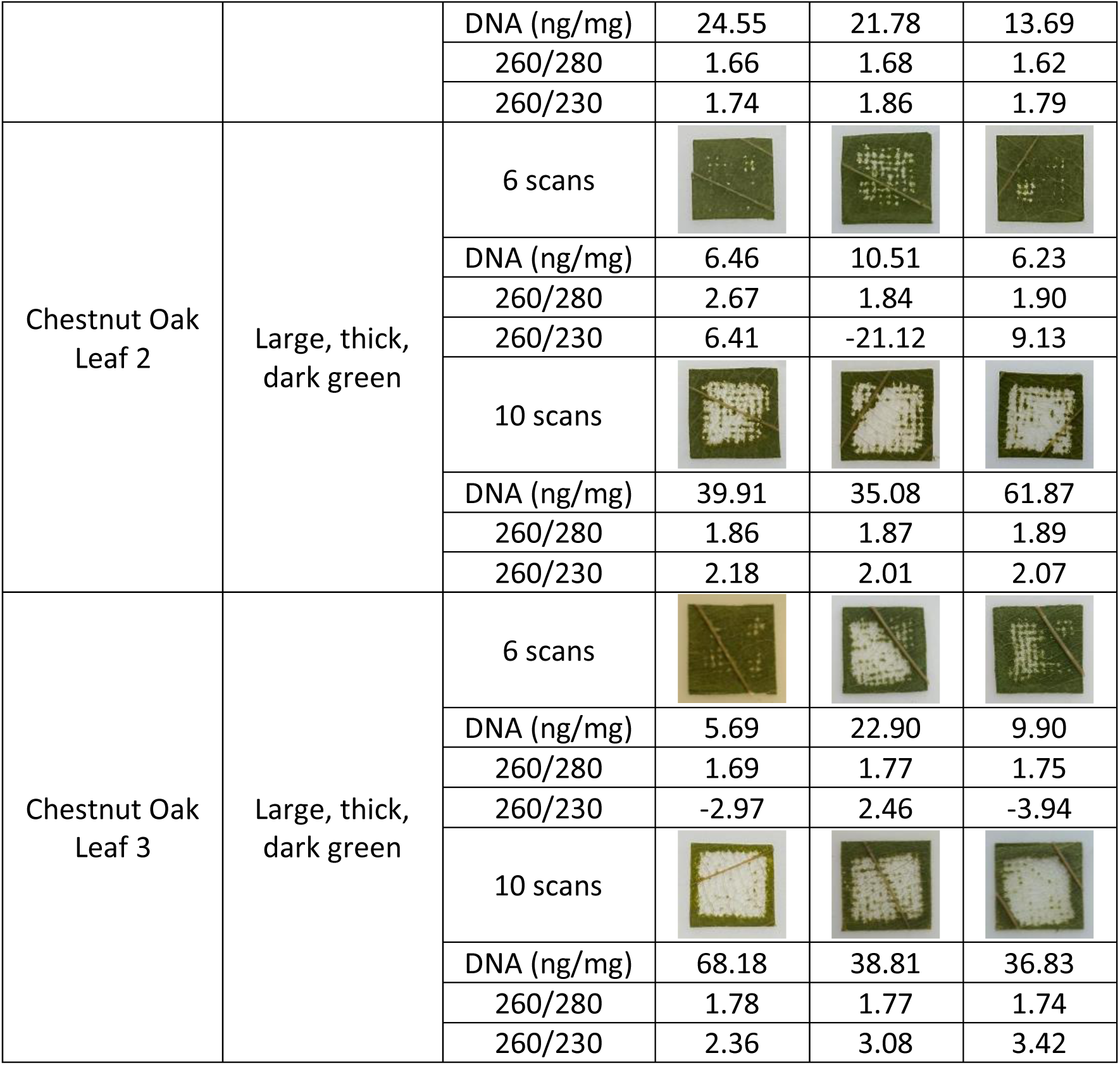
Chestnut oak tissue breakdown after FUSE processing demonstrates that greater tissue disintegration increases the concentration of DNA release. DNA quantification measurements are reported from Qubit^TM^ Fluorometer measurements, and 260/280 and 260/230 ratios are reported from Nanodrop^TM^ measurements.

| Leaf Species | Description | Data | Sample 1 | Sample 2 | Sample 3 |
| --- | --- | --- | --- | --- | --- |
| Chestnut Oak Leaf 1 | Large, thick, dark green | 6 scans |  |  |  |
|  |  | DNA (ng/mg) | 10.37 | 12.90 | 10.92 |
|  |  | 260/280 | 1.60 | 1.69 | 1.64 |
|  |  | 260/230 | 3.63 | 2.79 | 5.13 |
|  |  | 10 scans |  |  |  |
|  |  | DNA (ng/mg) | 24.55 | 21.78 | 13.69 |
|  |  | 260/280 | 1.66 | 1.68 | 1.62 |
|  |  | 260/230 | 1.74 | 1.86 | 1.79 |
| Chestnut Oak<br>Leaf 2 | Large, thick,<br>dark green | 6 scans |  |  |  |
|  |  | DNA (ng/mg) | 6.46 | 10.51 | 6.23 |
|  |  | 260/280 | 2.67 | 1.84 | 1.90 |
|  |  | 260/230 | 6.41 | -21.12 | 9.13 |
|  |  | 10 scans |  |  |  |
|  |  | DNA (ng/mg) | 39.91 | 35.08 | 61.87 |
|  |  | 260/280 | 1.86 | 1.87 | 1.89 |
|  |  | 260/230 | 2.18 | 2.01 | 2.07 |
| Chestnut Oak<br>Leaf 3 | Large, thick,<br>dark green | 6 scans |  |  |  |
|  |  | DNA (ng/mg) | 5.69 | 22.90 | 9.90 |
|  |  | 260/280 | 1.69 | 1.77 | 1.75 |
|  |  | 260/230 | -2.97 | 2.46 | -3.94 |
|  |  | 10 scans |  |  |  |
|  |  | DNA (ng/mg) | 68.18 | 38.81 | 36.83 |
|  |  | 260/280 | 1.78 | 1.77 | 1.74 |
|  |  | 260/230 | 2.36 | 3.08 | 3.42 |

The DNA extraction results show that leaf species influenced the DNA yield. Since the leaves chosen represented a range of angiosperm taxonomic diversity, it was expected that differences in physical and chemical properties would affect the quantity of DNA released. The control DNA concentration data was examined to determine the effect of species on DNA release. The American chestnut DNA yield was significantly lower than the average DNA yield from the control samples. The tulip poplar DNA yield was significantly greater than the average DNA yield from the control samples. The American chestnut leaves were the only samples described as brown-green, while the tulip poplar leaves were the only samples characterized as yellow-green. These sample characteristics suggest that the American chestnut samples were more mature than the tulip poplar samples at the time of collection [46]. It is common for older leaves to have greater amounts of secondary metabolites, which often cause low yield and poor quality DNA [14]. Therefore, it is likely that the age of the sampled leaves influenced inconsistencies in the quantity of the released DNA. As mentioned previously, it is also plausible that species-specific differences influenced the DNA yield. Further investigation into the physical and chemical properties of each leaf species would be needed to confirm this possibility.

The 260/280 and 260/230 ratios were measured to assess the quality of the DNA extracted with FUSE and conventional methods (Table 5). For American chestnut and chestnut oak, the 260/280 ratio was significantly higher for samples processed with six FUSE scans than for control samples. The 260/280 ratio for six FUSE scans was significantly lower than conventional methods for red maple leaves. No discernible trends were observed in 260/230 ratios, with values in expected norms for leaf tissue. However, the 260/30 ratio was significantly higher after ten FUSE scans than controls for chestnut oak leaves. This significant difference is likely the result of increased DNA release due to increased tissue breakdown. Some species processed with FUSE showed high standard error in 260/230 ratios. This result is likely due to incomplete tissue disintegration for some samples within these groups. However, it may also suggest that DNA extraction with FUSE affects the breakdown of compounds that contribute to turbidity in a purified sample.

**Table 5:** The quality of DNA released by FUSE is comparable to DNA released by conventional methods. FUSE requires less processing time than conventional methods and releases greater quantities of DNA. DNA quantification measurements are reported from Qubit^TM^ Fluorometer measurements, and 260/280 and 260/230 ratios are reported from Nanodrop^TM^ measurements.

| Sample Type | FUSE |  |  |  | Conventional Method |  |  |  |
| --- | --- | --- | --- | --- | --- | --- | --- | --- |
|  | Time (mm:ss) | DNA (ng/mg) | 260/280 | 260/230 | Time (mm:ss) | DNA (ng/mg) | 260/280 | 260/230 |
| American chestnut | 9:00 | 24.33 ± 6.51 | 2.34 ± 0.31 | 2.68 ± 3.51 | 30:00 | 6.22 ± 0.87 | 1.45 ± 0.01 | 0.52 ± 0.01 |
| Tulip poplar | 9:00 | 32.61 ± 7.82 | 1.77 ± 0.05 | 1.41 ± 0.34 | 30:00 | 28.37 ± 1.78 | 1.83 ± 0.01 | 1.99 ± 0.06 |
| Red maple | 9:00 | 35.32 ± 9.21 | 1.63 ± 0.04 | 1.95 ± 0.60 | 30:00 | 11.51 ± 1.95 | 1.85 ± 0.02 | 2.53 ± 0.19 |
| Chestnut oak<br>6 scans | 9:00 | 10.65 ± 1.74 | 1.84 ± 0.11 | 0.17 ± 3.00 | 30:00 | 17.17 ± 1.98 | 1.52 ± 0.01 | 0.66 ± 0.02 |
| Chestnut oak<br>10 scans | 15:00 | 37.85 ± 5.93 | 1.76 ± 0.03 | 2.28 ± 0.20 | 30:00 | 17.17 ± 1.98 | 1.52 ± 0.01 | 0.66 ± 0.02 |

The FUSE protocol used in this work involves a non-thermal tissue lysis process that has been shown to reduce the time required for DNA release. The acoustic parameters used in this study, particularly the PRF of 500 Hz, were chosen for this initial feasibility study based on preliminary tissue breakdown experiments. In previous work, Atlantic salmon muscle tissue was processed with FUSE using 10,000 pulses delivered at 25 Hz, which resulted in a total processing time of 6 minutes and 40 seconds [20]. In this study, 270,000 pulses were applied at 500 Hz to complete 6 FUSE scans, and 450,000 pulses were applied at 500 Hz to complete 10 FUSE scans, resulting in total processing times of 9 and 15 minutes. At higher PRFs, the time efficiency of FUSE was improved without inducing thermal effects. However, further increasing the PRF is likely to result in cavitation shielding effects that lower the effectiveness of each pulse [44, 45]. Future work will be necessary to explore the optimal pulsing parameters for the implementation of higher PRFs. We expect that implementing FUSE at higher PRFs will expand the applicability of FUSE to more robust tissue types and significantly decrease the tissue processing time.

### 3.3 DNA amplification

American chestnut samples were selected for amplification and sequencing. All FUSE and control samples were amplified with PCR following restriction digestion and adaptor ligation. The samples processed with FUSE and conventional methods amplified successfully, demonstrating that FUSE yielded high-quality DNA from leaf tissue suitable for PCR amplification. A subset of the amplified American chestnut libraries was visualized on a 2100 Bioanalyzer to examine the distribution of DNA fragment sizes for samples processed with FUSE and conventional methods (Figure 7). Most fragments for both the FUSE and control samples were within the expected range of 200-600 bp (Figure S2), suggesting that FUSE does not cause greater DNA fragmentation than conventional extraction methods.

**Figure 7:**
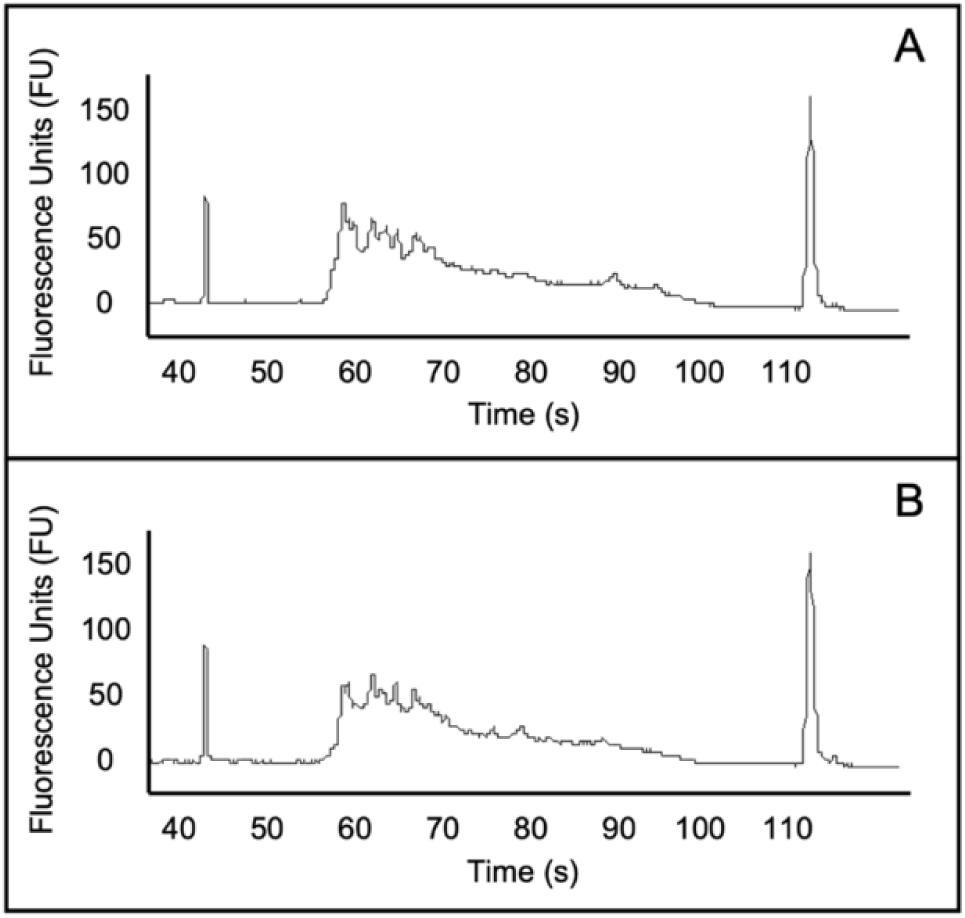
The distribution of DNA fragment sizes for an American chestnut sample processed with conventional methods (A) is comparable to the DNA fragment size distribution for a sample processed with FUSE (B). This result confirms that the integrity of DNA provided by FUSE is suitable for PCR amplification.

### 3.4 Sequencing

All American chestnut samples processed with FUSE and conventional methods provided high- quality next-generation sequencing reads (Accession Number: PRJNA837224). Because downstream applications of this technology are expected to be focused on identifying genetic variants from sequence data for population genetics and systematics, we estimated read depth for variable sites, which showed that FUSE samples had a depth comparable to controls. Read depth was moderately correlated between the two extraction methods (Figure S3), and mean depth was not significantly different (25.7 for FUSE and 27.1 for controls; P=0.155 based on a Wilcoxon rank-sum test). Among individual FUSE and control samples, read depth was fairly consistent for those processed with FUSE and conventional methods, suggesting that the FUSE protocol yields DNA with quality comparable to conventional methods, and the DNA is suitable for NGS analyses (Figure 8).

**Figure 8:**
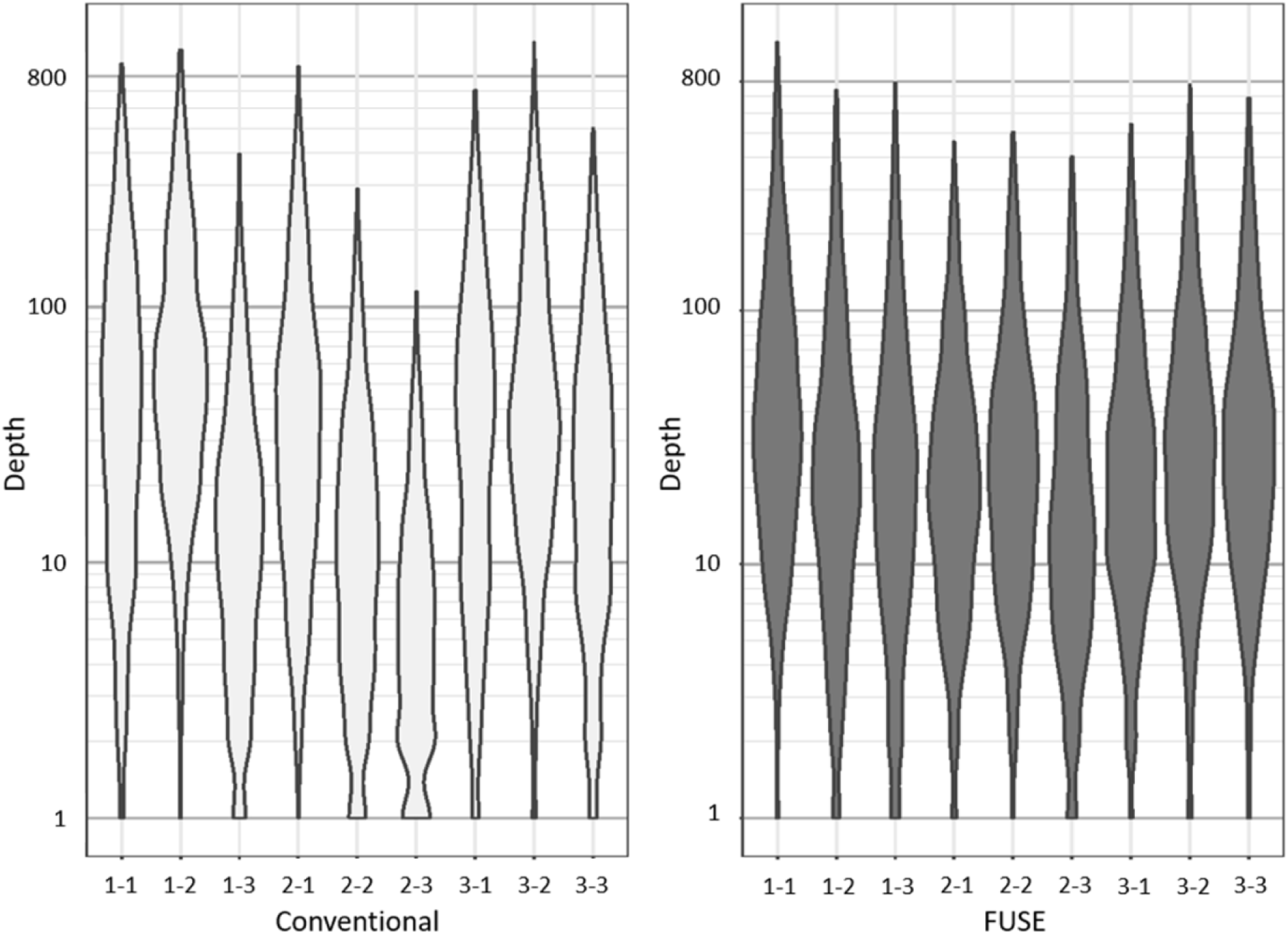
FUSE provides DNA suitable for next-generation sequencing. The uniformity of read depth across conventional and FUSE samples is comparable. The x-axis labeling represents the leaf and sample number, such that 1-2 identifies the read depth for leaf 1, sample 2.

## 4. Conclusion

This study assessed the efficacy of our recently developed FUSE protocol in plant tissues by testing samples from American chestnut, tulip poplar, red maple, and chestnut oak leaves. The success of the FUSE protocol was determined by visualizing the extent of tissue breakdown observed after FUSE sonication, measuring the quantity and quality of the released DNA, and evaluating the suitability of DNA extracts for genetic analyses. PCR amplification and NGS were done to assess the utility of the released DNA in genomic workflows. In accordance with previous work that established the effectiveness of FUSE for releasing DNA from Atlantic salmon muscle tissue [20], the results of this study demonstrate that FUSE can provide high quantities of DNA suitable for amplification and sequencing in less time than conventional plant extraction methods. Additionally, these results suggest that the input sample mass required by FUSE is less than what is necessary for conventional extraction methods, which could be advantageous in future work that aims to develop field-deployable FUSE systems for conservation efforts. Overall, this study shows that the applications of FUSE can be extended to plant tissue, a robust tissue that is more resistant to mechanical breakdown and has a chemical composition that has traditionally made DNA accessibility more challenging [4, 13, 14]. In conjunction with previous findings [20], these results suggest that FUSE could be used as a novel platform for DNA extraction capable of accelerating workflows for a variety of sample types.

## Acknowledgments

This work was funded by a grant from the Gordon and Betty Moore Foundation (Grant #8518). We would like to specifically thank Dr. Sara Bender and the Moore Foundation’s Science Program for their ongoing support of this project. Finally, we would like to thank Conservation X Labs, the National Geographic Society, the Virginia Tech Department of Biomedical Engineering and Mechanics, and the Virginia Tech Institute for Critical Technology and Applied Science for their support of this work.

## Data Accessibility Statement

All genetic sequences generated in this work are available on Genbank. Next-generation sequencing data can be accessed with the Accession number: PRJNA837224.

