## Supplementary figures and images for "DNA Release from Complex Plant Tissue using Focused Ultrasound Extraction (FUSE)"

### Figure S1

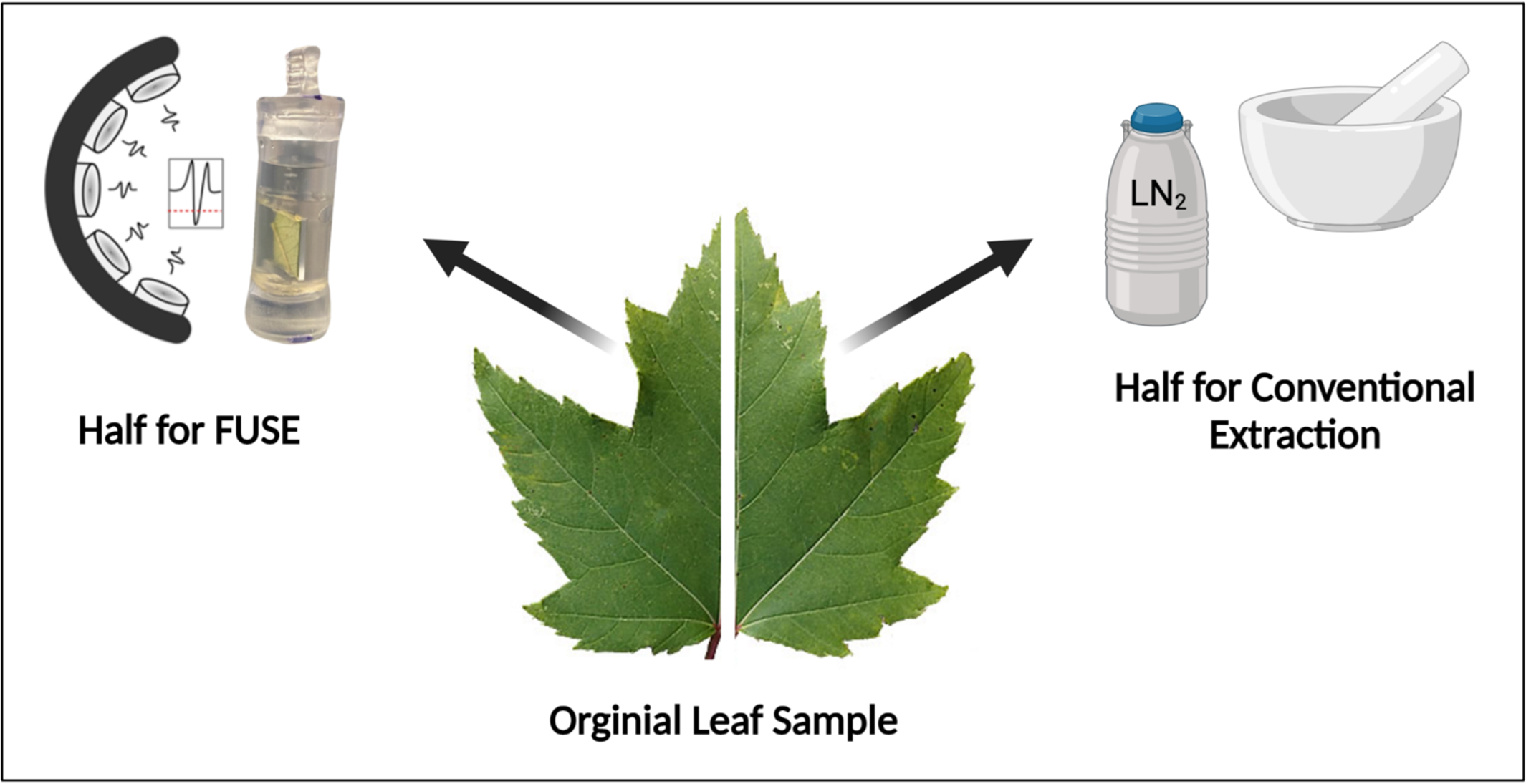

### Figure S2

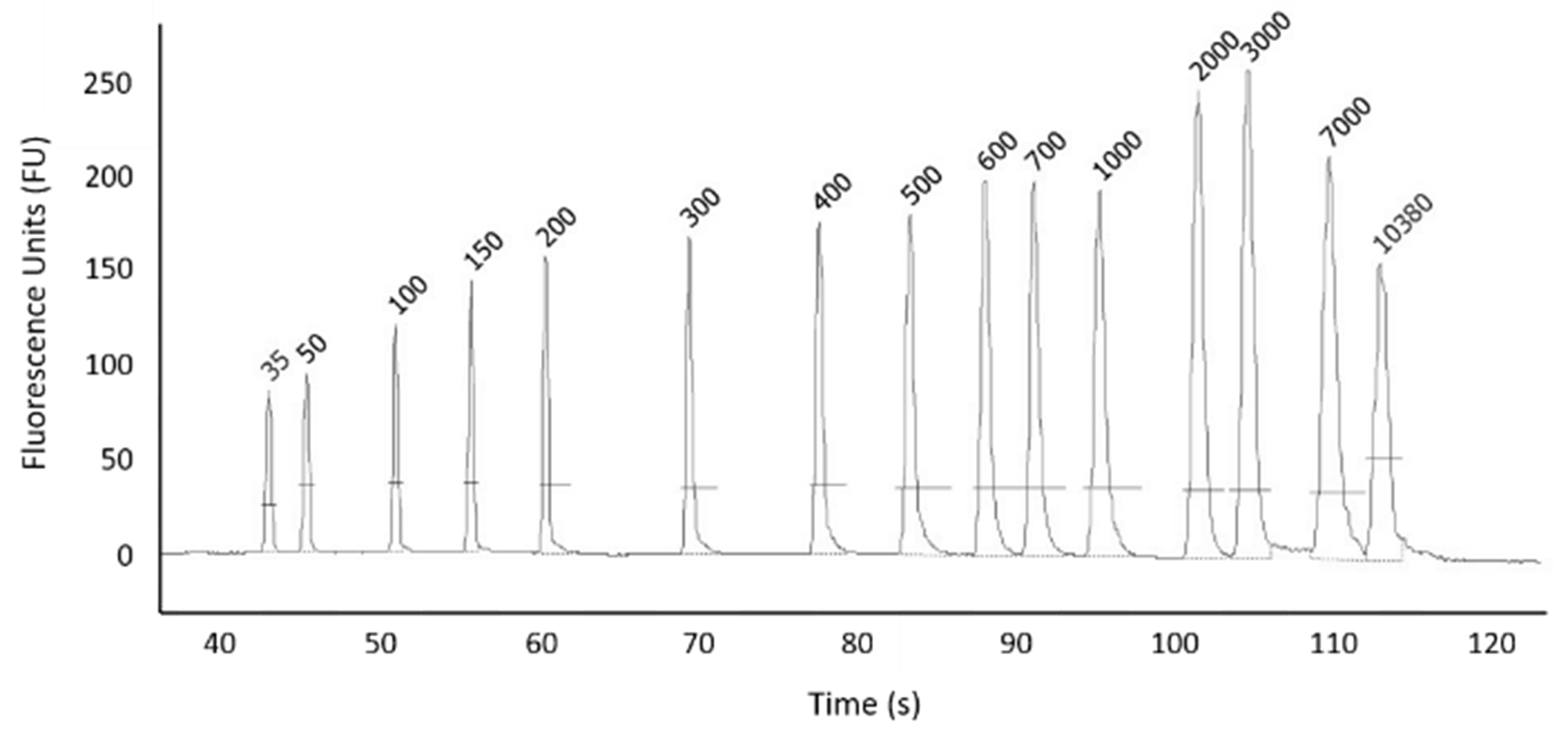

### Figure S3

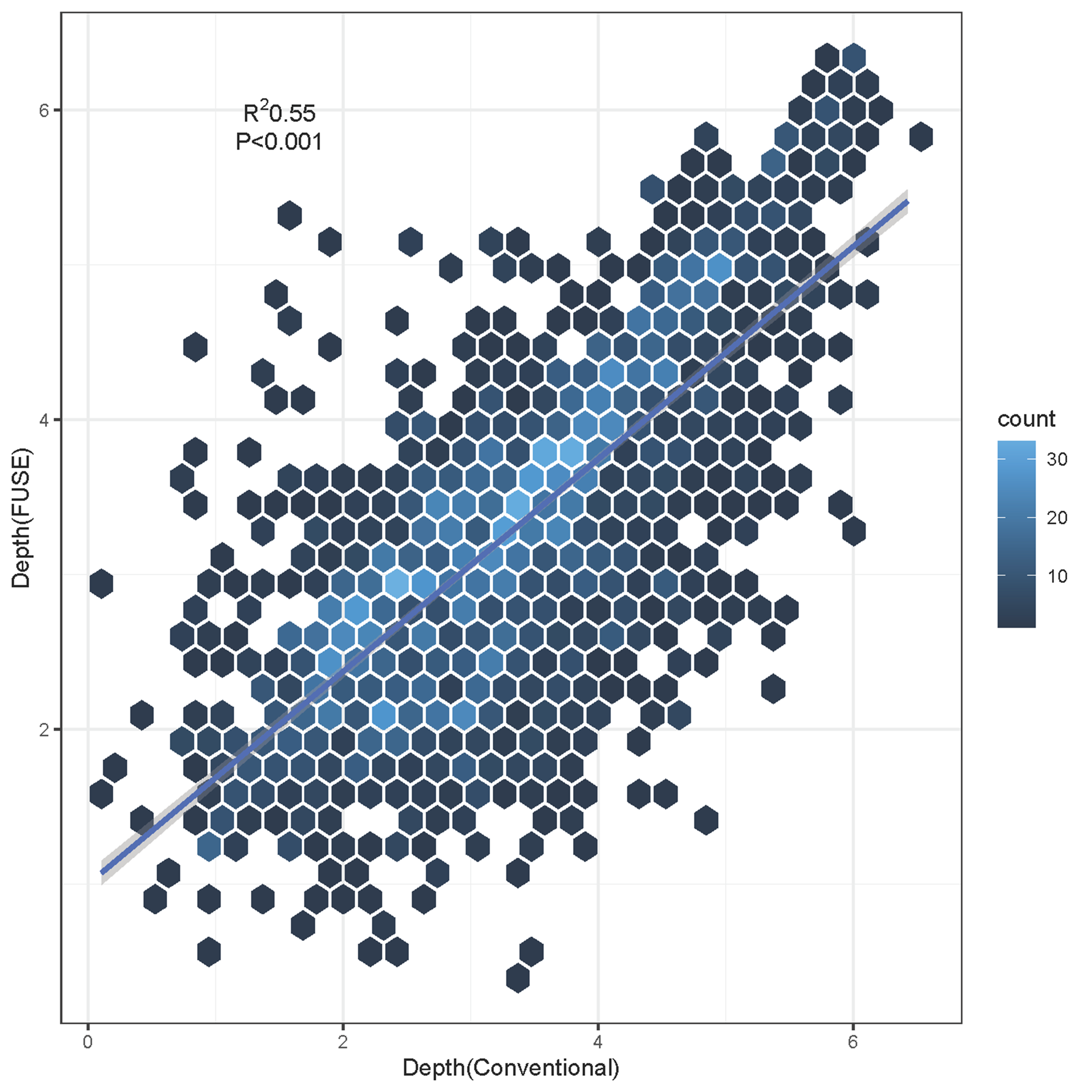
